# Urban green space exposure reduces subjective stress and physiological arousal

**DOI:** 10.64898/2026.05.18.724862

**Authors:** Dilber Korkmaz, Qingyue Bi, Michelle Moller, Julian Koenig, Jan Peters

## Abstract

Stress is a major risk factor for mental disorders, and urban living is a key environmental contributor. Nature exposure may promote stress recovery and mental health, but how physiological arousal and subjective stress change across green versus gray space during naturalistic urban mobility is poorly understood. This preregistered study (https://doi.org/10.17605/OSF.IO/HF4RW) employed geolocation-based ambulatory assessment to examine psychophysiological arousal and subjective stress during transitions between urban green and gray environments. Thirty-six healthy urban residents completed a counterbalanced circular walking route in Cologne, Germany, with continuous GPS, cardiovascular, and electrodermal recording alongside ecological momentary assessment of subjective stress, affect, and exertion. Green compared to gray spaces were associated with lower subjective stress and higher affective well-being, with cardiac indices reflecting reduced autonomic arousal during green space exposure. Autonomic changes surrounding environmental transitions persisted beyond the immediate transition window, suggesting that physiological benefits of green space exposure extend into subsequent gray environments. These findings underscore the public health potential of urban green infrastructure for preventing stress-related mental health conditions.

## Introduction

Stress-related mental health conditions represent a growing global health burden, with depression and anxiety among the most prevalent disorders worldwide (1). Stress involves the coordinated activation of physiological and psychological systems in response to environmental demands that challenge or threaten an individual’s well-being (2,3). The autonomic nervous system (ANS) and the hypothalamic-pituitary-adrenal axis (HPA) play central roles in mediating physiological stress responses, including increases in heart rate (HR), cortisol release, and altered autonomic regulation (2,4–6). Sympathetic activation produces rapid increases in HR and electrodermal activity (EDA), whereas parasympathetic activity is reflected in elevated heart rate variability (e.g., root mean square of successive differences, RMSSD), an index of vagally mediated cardiac regulation (7–13). ANS indices offer complementary windows into physiological arousal, capturing sympathetic activation and parasympathetic recovery across different timescales. At the psychological level, stress manifests as negative affect, emotional distress, cognitive strain, and impaired well-being in response to demands perceived as exceeding one’s coping resources (14,15). When stressors persist or exceed coping capacity, these responses contribute to the development and maintenance of mental health conditions, with chronic environmental stress consistently identified as a risk factor for mental disorders (3,16).

Urbanization has emerged as a significant environmental driver of stress and stress-related mental health conditions (17,18). Urban residents show consistently higher rates of mental disorders compared to rural populations, a pattern documented across diverse geographic and cultural contexts (19–21). This elevated risk may be partly attributable to chronic exposure to urban stressors, including noise, overcrowding, climate change, and air pollution, with environmental exposures directly influencing neuroendocrine, inflammatory, and autonomic pathways contributing to depression and anxiety (19,22,23). As urban populations continue to grow globally, identifying accessible and scalable strategies to reduce the stress burden of urban living has become an important public health priority.

Nature and green space exposure have been proposed as one such strategy, grounded in two theoretical frameworks. *Stress Recovery Theory* draws on psychoevolutionary principles to argue that natural environments elicit positive affective responses and reduce sympathetic nervous system activation, enabling rapid psychophysiological recovery from stress (24). *Attention Restoration Theory* offers a complementary cognitive perspective, proposing that natural settings engage involuntary attention in an effortless manner, allowing the directed attentional resources depleted by urban cognitive demands to replenish, thereby reducing mental fatigue and supporting emotional well-being (25). Together, these frameworks predict that green environments should reduce stress across psychological and physiological levels. Since these levels may not always align (26–29), comprehensive assessment requires simultaneous measurement at both (30,31).

Existing evidence spans controlled laboratory and real-world field settings. Laboratory studies using nature videos or images have shown that viewing natural compared to urban scenes accelerates physiological stress recovery through faster normalization of HR and skin conductance and increases in vagally mediated heart rate variability, indicating a shift toward parasympathetic dominance (24,32–34). Field-based studies extend these findings to naturalistic settings, with walking in natural versus urban environments is associated with reduced rumination, improved mood, and reduced physiological arousal (35–39). Meta-analytic and epidemiological evidence consistently links green space exposure to psychophysiological stress reduction and lower rates of depression, anxiety, and common psychiatric disorders across diverse populations (40–47). However, laboratory paradigms lack ecological validity, as passive exposure to simulated environments does not capture the dynamic and multisensory nature of real-world urban mobility. Existing field studies have predominantly operationalized green space as a static between-condition variable, and have rarely integrated continuous physiological monitoring with comprehensive self-report and real-time GPS-verified environmental exposure, limiting the extent to which the joint dynamics of subjective and physiological measures can be examined in situ.

Ambulatory assessment methods enable simultaneous continuous physiological recording and self-report with high temporal resolution and ecological validity (48–50). Ecological momentary assessment (EMA) captures subjective stress as it unfolds across changing environments rather than relying on retrospective recall (51). Characterizing stress across repeated environmental transitions is particularly relevant given that cumulative daily encounters with urban green space may be more consequential for mental health than single isolated exposures. Yet most existing research has focused on forests or decentralized parks rather than the green spaces embedded in the everyday urban environment that residents regularly encounter.

The present preregistered study employed geolocation-based ambulatory assessment while healthy urban residents completed a circular route involving repeated transitions between green and gray urban environments in Cologne, Germany. Subjective stress, affective well-being, and physical exertion were assessed via EMA, while HR, heart rate variability, and EDA were recorded continuously. GPS tracking verified environmental exposure and transition timing. We preregistered the hypotheses that green compared to gray space exposure would be associated with reduced subjective stress, improved affective well-being, and reduced psychophysiological arousal.

## Method

### Sample

Thirty-six healthy participants (*n* = 8 male, *M_age_* = 21.97, *SD =* 2.88) were included in the final sample. Data were initially collected from *N* = 54 participants; *n* = 18 participants were excluded due to signal artefacts (*n* = 5), electrode disconnections (*n* = 3), or excessive movement artefacts (*n* = 10). One participant did not provide usable GPS data; however, this participant was retained in the final sample as environmental transition timepoints were systematically recorded in real time by the experimenter during the walk and used in place of GPS-derived timestamps. Furthermore, three participants in the final sample (*N* = 36) showed a single missing EMA prompt during specific segments of the route: Baseline (*n* = 2), Gray (*n* = 1), and Post (*n* = 1). These participants were retained for all analyses, as their remaining subjective responses provided sufficient data for segment-level comparisons. The final sample size of *N* = 36 is consistent with our a priori power analysis conducted using G*Power (52). Based on an average effect size of *d* = .51 reported by Gong et al. (2024) for physiological reactivity contrasts between urban green and gray space exposure, a one-tailed paired t-test with α = .05 and power (1-*β*) = .80 yielded a required sample size of *N* = 27. The target sample was set to account for anticipated data loss due to artefacts and technical issues inherent to ambulatory physiological recordings. In the preregistration we aimed for a gender-balanced sample, but due to time constraints during data acquisition this could not be fulfilled. Inclusion criteria comprised no history of cardiovascular, psychiatric, or neurological disorder, and no current medication use or drug abuse. All participants provided written informed consent and received course credit points as reimbursement. The study was approved by the ethics committee of the University of Cologne (code: DKHF0251) and conducted in accordance with the Declaration of Helsinki. Data were collected between October 2024 and February 2025 in Cologne, Germany.

### General Procedure

This study was preregistered and is available at OSF (https://doi.org/10.17605/OSF.IO/HF4RW). In a single-day within-subjects design, participants navigated a predetermined circular urban walking route (*M*_distance_ = 4.02 km, *SD* = 0.06; *M*_duration_ = 61.82 min, *SD* = 3.62) transitioning between green and gray environments while undergoing ambulatory assessment of EMA, GPS, and physiological data. Environmental exposure order was counterbalanced across participants via alternating start points, with both groups walking the same route in opposite directions following the baseline segment. Data collection was conducted during daylight hours (9 AM–5 PM) and postponed under extreme weather conditions (e.g., thunderstorms, heavy rain, or high winds).

Prior to the walk, participants provided written informed consent and completed questionnaires assessing socio-demographic characteristics, stress, and personality in the laboratory (see *supplementary methods*). They then received a visual overview of the route, and a trained researcher attached mobile ECG and EDA sensors. Participants were equipped with a study smartphone for GPS tracking and EMA delivery, and instructed to refrain from conversing with others, listening to music, or using their private phone during the walk. Participants walked alone but followed the experimenter at approximately 20 meters distance, enabling real-time notation of route deviations, unexpected disruptions, and environmental transition timestamps for GPS verification. Participants were prompted to respond to six EMA probes assessing momentary subjective stress, physical exertion, and affective well-being on visual analogue scales (VAS). Upon return, sensors were removed and participants reported any noteworthy events experienced during the walk.

### Ambulatory assessment

#### GPS-based location tracking and route description

Participants were equipped with a study smartphone for continuous GPS location tracking using the GPS Logger App (version 2.1.12) for Android, recording instantaneous position at one-second intervals to ensure high spatial accuracy. Time and date were synchronized with the mobile physiological sensors via the linked computer to enable precise temporal alignment of location and physiological data. The walking route comprised six sequential segments: a baseline segment, two green segments, two gray segments, and a post-exposure segment. The baseline and post-exposure segments consisted of urban built environment and served as the start and end points of the route, leading away from and returning to the university campus. The green and gray segments were selected to represent distinctly contrasting urban environments. In the green segments participants traversed the Hiroshima-Nagasaki-Park in Cologne, a publicly accessible urban green space characterized by a central pond, tree-lined paths, open meadows, and wildlife including waterfowl and small mammals. The area is largely shielded from motorized traffic and dominated by natural visual stimuli. The gray segments followed busy urban arterial roads characterized by multi-lane motorized traffic, dense commercial and residential building facades, paved surfaces, and minimal vegetation (see Fig. 1).

**Figure 1.**
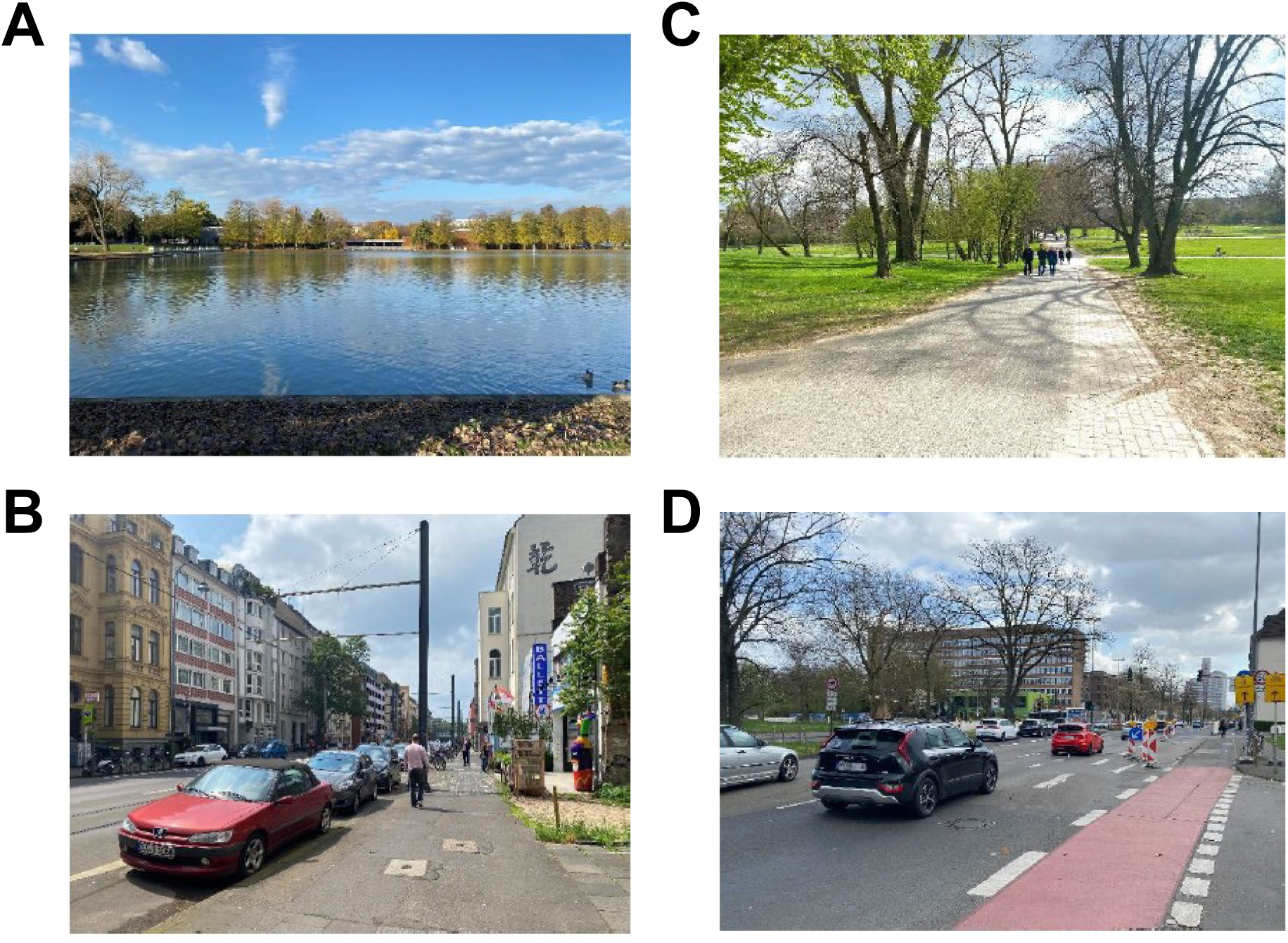
Representative visual pictures of the green and gray urban segments traversed during the study in Cologne, Germany. Panels A and B depict typical scenes from the green segments and was photographed on the path near the central pond in the Hiroshima-Nagasaki-Park; panel B shows the within the vegetation-dominated segment of the same park, featuring slight slopes visible in both directions and. Panels C and D depict typical scenes from the gray segments and was photographed on the heavily trafficked urban road characterized by building facades with paved surfaces and minimal vegetation. All photographs were taken during the data collection period and are representative of the environmental conditions encountered by participants.

#### Ecological momentary assessment

Subjective responses were assessed using single-item VAS measures capturing momentary stress level (“*How stressed do you feel right now?*“; 0 = not at all to 100 = very stressed), affective well-being (“*How do you feel right now?*“; 0 = very unwell to 100 = very well), and perceived physical exertion (“*How physically exerted do you feel right now?*“; 0 = not at all to 100 = very exerted). Participants were instructed to report their current, momentary state at the time of each prompt. EMA prompts were delivered via the study smartphone through an auditory notification and vibration, initiated by the experimenter at predefined locations along the route. Upon receiving a prompt, participants were instructed to stop walking and stand at a safe position at the side of the path before completing the assessment, such that all responses were provided while stationary. Participants completed the EMA a total of six times.

#### Physiological recording

Cardiovascular activity and EDA were recorded using the EcgMove4 and EdaMove4 devices (Moviesense ©, 53), enabling continuous acquisition of ECG and EDA signals. Data were stored according to the system time of the paired computer, ensuring synchronized timestamps across physiological channels. The ECGmove4 sensor was attached to the participant’s left chest using disposable, pre-gelled Ag/AgCl electrodes. Raw single-channel ECG data were recorded at a sampling rate of 1024 Hz, with HR (bpm) and heart rate variability (as indexed as RMSSD in ms) extracted as the primary cardiac indices. The EdaMove4 was attached to the non-dominant wrist, with electrodes placed on the thenar and hypothenar eminences of the non-dominant palm. A constant DC voltage of 0.5 V was applied to assess skin conductance, sampled at 32 Hz, from which skin conductance level (SCL, µS), skin conductance response peaks (SCR peaks in count), and mean SCR amplitude (µS) were derived as indices of sympathetic arousal.

### Data analysis

#### GPS-based location tracking processing

GPS data were continuously recorded during the route, capturing latitude, longitude, elevation, speed, and timestamps at one-second intervals. Raw data were exported in GPX format and processed in R (version 4.2.2, 54). Route distances between transition points were computed using the Haversine formula implemented in the *geosphere* package (55). Timestamps were converted to local time using *lubridate* (56), to ensure temporal consistency across participants and mobile sensors. Interactive route maps were generated using the *leaflet* package (57,58). Sequential transition points along the route were labeled T0 through T6, defined *a priori* based on specific landmarks along the route for each environmental segment. A custom R script was developed to automatically match recorded GPS points to the predefined target coordinates for each participant, while preserving the sequential order of transitions. Given the expected positional uncertainty of smartphone-based GPS apps in dense urban environments, arising from satellite geometry, signal multipath, and building reflections (59), a buffer radius of 20 meters was applied. For all participants, the dataset contains the GPS coordinate target transition points (T0-T6), matched nearest GPS coordinates, timestamp, and geodesic distance between the target and the matched coordinate for each transition point.

Segments were defined as the intervals between consecutive transitions, yielding six consecutive segments per participant: a baseline period (T0–T1), four alternating environmental exposure segments corresponding to green and gray conditions (T1-T2, T2-T3, T3-T4, T4-T5), and a post-exposure period (T5–T6). Segment duration was calculated as the elapsed time between transitions, and segment distance as the cumulative geodesic distance between successive GPS points using the Haversine formula. Average speed was derived from instantaneous speed values recorded at each GPS point and averaged across the segment. The resulting segment-level metrics and transition points were used for subsequent statistical analyses.

#### Ecological momentary assessment

EMA data comprised six measurement points administered across the experiment: Baseline, Green 1/Gray 1, Green 2/Gray 2, and Post. For each participant, mean ratings of perceived stress, affective well-being, and physical exertion were calculated separately for green and gray environments. To obtain a single estimate per environment per participant, responses from repeated exposures to the same environmental condition were averaged. As preregistered, a repeated measures ANOVA was first conducted to confirm the absence of significant differences between the first and second exposure within each environment (e.g., Green 1 vs. Green 2), thereby validating the aggregation approach. Green and gray environments were then compared using preregistered paired-samples t-tests based on participant-level means, with Cohen’s *d* reported as the effect size for dependent samples. To account for the hierarchical and repeated-measures structure of the data, a preregistered linear mixed-effects model (LMM) was additionally fitted to the full, non-aggregated EMA dataset, serving as the primary inferential approach. For each outcome variable, fixed effects included environment (Green vs. Gray), measurement number (first vs. second exposure within each environment), and order group (Green-first vs. Gray-first). Note that the inclusion of measurement number and order group as fixed effects represents an extension of the preregistered model specification to more fully account for the counterbalanced design.

#### Physiological data

Raw physiological data were exported from the Movisens © devices in binary format. All preprocessing and feature extraction were performed in Python (version 3.11) using NeuroKit2 (60). ECG signals were converted to microvolts according to manufacturer specifications, cleaned to remove noise and baseline drift, and R-peaks were automatically detected. Consecutive RR intervals were calculated in milliseconds. EDA signals were filtered to remove high-frequency noise and decomposed into tonic and phasic components to extract SCL, SCR peaks, and SCR amplitude. Each measurement was timestamped against the absolute recording start time, enabling precise temporal alignment with GPS-derived transition points. Physiological data were aligned to GPS-defined segments representing the intervals between consecutive transition points (T0–T6), comprising a baseline period (T0–T1), four alternating environmental exposure segments corresponding to urban green and gray conditions, and a post-exposure period (T5–T6). ECG artifact rejection was applied in two stages: RR intervals outside 0.3–1.5 s (40–200 BPM) were removed, followed by exclusion of beats where successive RR intervals deviated by more than 20% from the preceding interval (61). Mean HR (bpm) and heart rate variability (indexed as RMSSD in ms) were computed across the full duration of each segment. EDA decomposition yielded mean SCL, mean SCR amplitude, and SCR peaks per segment; the latter was derived by averaging per-bin peak rates across 60-second bins to provide a duration-robust estimate (10).

Two complementary analytical approaches were applied to the preprocessed physiological data. *Segment-based analyses* were preregistered, and examined sustained physiological differences between environmental conditions aggregated across the exposure segments. *Transition-based analyses* examined the dynamic changes in autonomic responses in 60-second time bins surrounding each environmental transition using a principal component analyses (PCA)-derived multivariate index of autonomic arousal. Transition-based analyses and the associated PCA were not preregistered and were conducted to examine the temporal dynamics of autonomic effects around environmental transition points, complementing the segment-level analyses.

#### Segment-based analyses

All physiological variables were standardized using within-subject z-scoring computed across all six segments per participant, removing between-subject differences in absolute physiological levels while preserving within-person variation. The two green and two gray exposure segments were then averaged per participant to yield a single green and gray score per physiological variable. Within-subject differences were assessed using paired-samples t-tests with Cohen’s *d* as the effect size measure. To control for multiple comparisons across the five physiological variables, Bonferroni correction was applied, yielding an adjusted significance threshold of α = 0.01. A multivariate autonomic index was additionally derived via PCA on the z-scored values of all five physiological variables across the four experimental segments, treating each participant × segment as an observation. PC scores were extracted per participant per segment, averaged within each condition (e.g., Green 1 and Green 2), and compared using paired-samples t-tests.

One green segment was identified *post hoc* as a potential confound based on the presence of a notable relatively steep incline slope in this segment that was overlooked during the initial planning of the route. A paired-samples t-test comparing HR between the affected and unaffected green segments revealed significantly higher HR in the segment containing the slope (*M* = 102.31, *SD* = 11.27) compared to the unaffected segment (*M* = 98.00, *SD* = 11.10) with *t*(35) = 7.36, *p* < .001. This is consistent with effects related to sustained effort from the slope, rather than environmental stress responding. To address this issue, all analyses of physiological measures were repeated with the slope segment removed. For full transparency, we report results for the full data set and the slope-corrected data set in the main paper. The slope-affected segment corresponded to the second green exposure for green-first participants and the first green exposure for gray-first participants, ensuring that the exclusion was applied symmetrically across counterbalance groups.

#### Transition-based analysis

Transition-based analyses were conducted as a complement to the preregistered segment-based approach and were not preregistered. The isolated autonomic response to environmental transitions was examined by centering physiological time series on GPS-derived transition timestamps at segment boundaries T2, T3, and T4. Each transition timepoint was defined as the moment the participant crossed the environmental boundary. Three transitions were examined, which differed by counterbalance group: green-first participants experienced a green-to-gray transition at T2 and T4 and a gray-to-green transition at T3, while gray-first participants experienced a gray-to-green transition at T2 and T4 and a green-to-gray transition at T3.

Physiological time series were segmented into consecutive non-overlapping 60-second time bins, with the center bin 0 defined as the interval spanning 30 seconds before to 30 seconds after the transition. Preceding and following bins were indexed from -4 to +4, respectively. To ensure a consistent bin structure across participants, the shortest green or gray segment duration across the sample was identified and divided by 60 to determine the maximum number of complete 60-second bins consistently available for all participants. This value was then divided by two to establish the number of pre- and post-transition bins included per transition, with any remaining edge time discarded. Data were structured in long format with each row representing a single participant × bin × variable combination. Outliers were identified per participant per variable using z-score thresholding (|z| > 3) and replaced via linear interpolation between adjacent valid observations to preserve time-series continuity. Physiological variables were then standardized using within-subject z-scoring computed across all bins and transitions, following the same rationale as the segment-based analysis but applied to the bin-level time series rather than segment-level means.

Again, a PCA was conducted on the within-subject z-scored values of all five physiological variables across all participant × bin observations across all three transitions. PC scores were aggregated at the participant level ensuring each participant contributed at most one transition per direction. For green-first participants, the two green-to-gray transitions at T2 and T4 were averaged to yield a single green-to-gray score per bin, and the single gray-to-green transition at T3 was used directly. For gray-first participants, the two gray-to-green transitions at T2 and T4 were averaged to yield a single gray-to-green score per bin, and the single green-to-gray transition at T3 was used directly. Group-level means and standard errors were computed for visualization and inference.

One transition per counterbalance group was identified as potentially confounded by incline slope effects, corresponding to the transition immediately following the slope-affected green segment. Specifically, T4 (green-to-gray) was affected for green-first participants and T2 (gray-to-green) for gray-first participants. To isolate psychophysiological responses to environmental change from physical exertion effects, all transition analyses were conducted both including and excluding these transitions, with results reported accordingly.

## Data availability

The data that supports the findings of this study will be made available upon publication via the Open Science Framework.

## Results

### GPS

Segment-level GPS data are summarized in Table 1, with both green and gray segments combined in one single segment. Participants’ movement patterns differed across the segments. The green segments were covered at the highest average speed, followed by gray segments, with baseline and post segments showing the slowest speeds. Distances and durations reflected the predefined route structure, with the longest segments corresponding to environmental exposures in gray and green, and shorter segments observed during baseline and post phases. These findings confirm that participants navigated the planned route and that segment-level movement metrics were consistent across participants.

**Table 1.**
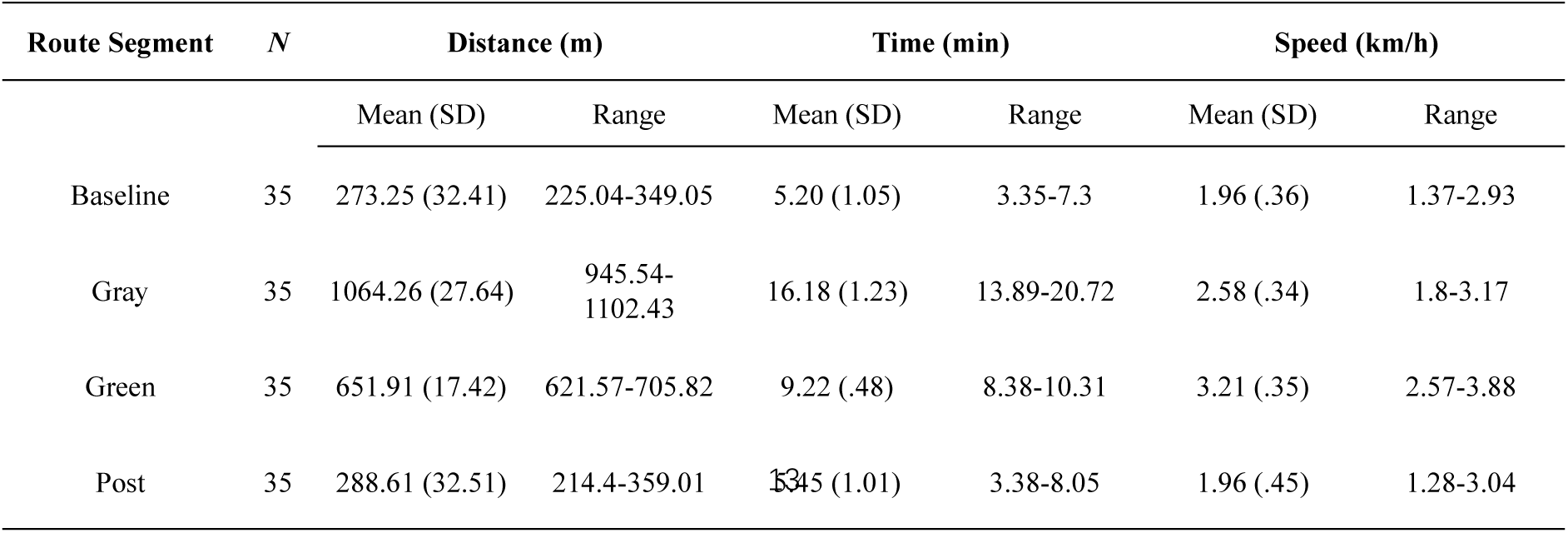
Descriptive statistics of GPS-derived route segment metrics across four phases of the urban walking route. Distance (m), time (min), and walking speed (km/h) are reported as mean (SD) and range per segment, aggregated across both order groups. GPS data were available for *N* = 35 participants, as one participant did not provide usable GPS data and experimenter-logged transition timepoints were used in place of GPS-derived timestamps for this participant; segment-level metrics could therefore not be computed. Green and Gray reflect the aggregated green and gray urban environment segments respectively. Baseline and Post reflect the start and end segments of the route.

GPS data were evaluated at the level of individual transition points (T0–T6). For each participant and each transition, the geodesic distance between the predefined target coordinate and the closest recorded GPS point was computed. Across both conditions, mean matching distances were small (*M*_green_ = 19.1 m, *SD* = 1.8; *M*_gray_ = 18.8 m, *SD* = 2.1), indicating that recorded GPS positions closely corresponded to the intended transition locations. All participants successfully completed the full walking route in both conditions, passing through all seven transition points (see Fig. 2). For the participant with unusable GPS data, the transition protocol recorded by the research assistant (timestamps) during the walk was used, with target coordinates assigned based on the predefined reference points for each transition. This procedure ensured that this participant could be retained in all subsequent transition-based analyses.

**Figure 2.**
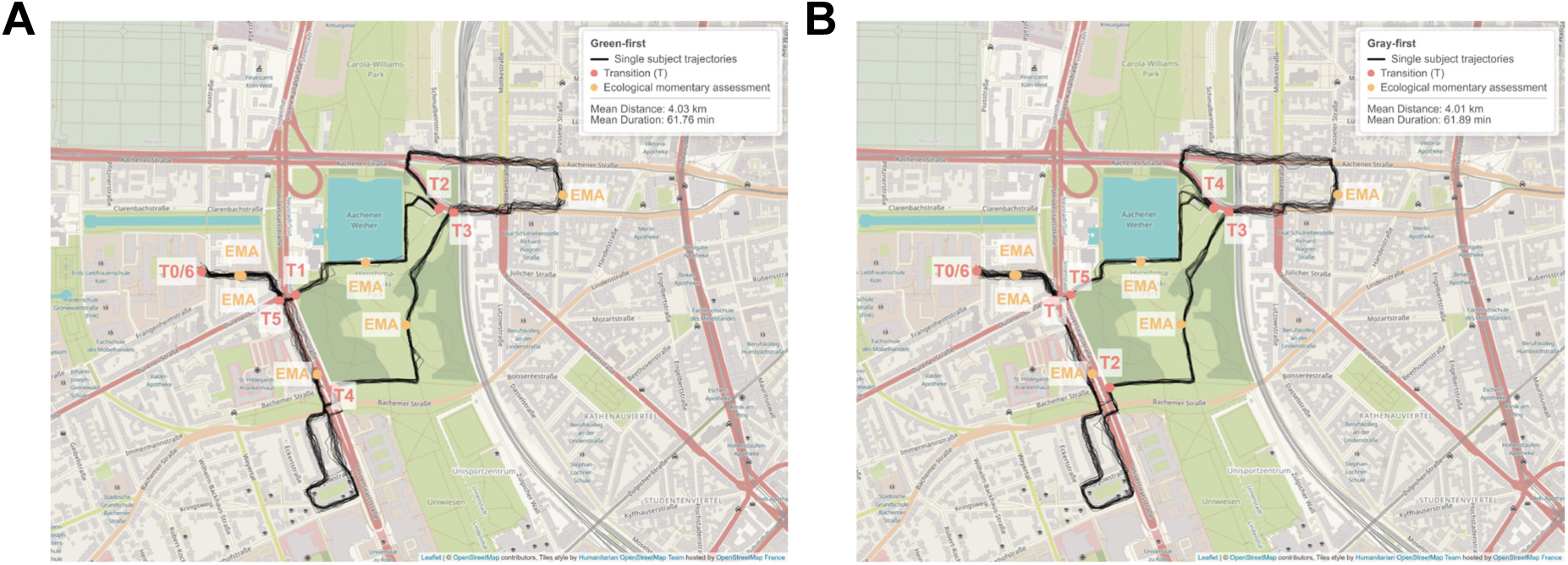
Illustration of the urban walking route in Cologne, Germany. Individual participant movement trajectories derived from GPS-based location tracking are depicted by black lines, separately for green-first (panel A, *n* = 18) and gray-first participants (panel B, *n* = 17). Red markers indicate the seven environmental transition points (T0–T6), with labels reflecting the order-specific environmental sequence for each group. Blue markers indicate a priori ecological momentary assessment measurement locations. Visualization created using Leaflet (https://leafletjs.com/), © OpenStreetMap contributors, CC-BY-SA (https://creativecommons.org/licenses/by-sa/2.0/), tiles style by Humanitarian OpenStreetMap Team (https://www.hotosm.org/) hosted by OpenStreetMap France (https://www.openstreetmap.fr/), implemented in RStudio with the leaflet package.

### Ecological momentary assessment

Prior to aggregating repeated exposures within each environment for the main analyses, a 2 (Environment: Green vs. Gray) × 2 (Time: first vs. second exposure) repeated measures ANOVA was conducted for each outcome variable to verify the equivalence of repeated exposures. No significant Environment × Time interactions were observed for any outcome (all *p* > .14), supporting the aggregation of first and second exposures within each environment. A significant main effect of Time was found for physical exertion (*F*(1, 35) = 8.60, *p* = .006), reflecting a general increase in perceived exertion across the route irrespective of environment. Descriptive trajectories and segment-level means for all six measurement points, depicted separately by order group, are provided in the *supplementary information* (see Fig. S1 and Table S1).

Within-subject comparisons between the green and gray environments were conducted using paired-samples t-tests on participant-level mean ratings, aggregating repeated exposures within each environment. Participants reported significantly lower stress in the green (*M* = 18.94, *SD* = 17.64) than in the gray environment (*M* = 26.03, *SD* = 19.76), *t*(35) = –3.97, *p* < .001, *d* = 0.66, and significantly higher affective well-being in green (*M* = 77.24, *SD* = 11.33) than in gray (*M* = 69.18, *SD* = 14.73), *t*(35) = 4.27, *p* < .001, *d* = 0.71. Physical exertion did not differ significantly between green (*M* = 20.24, *SD* = 10.66) and gray (*M* = 20.96, *SD* = 10.68), *t*(35) = –0.66, *p* = .52, *d* = 0.11. These comparisons are illustrated in Fig. 3. LMMs were used to examine the effects of environment, measurement number, and order group on stress, affective well-being, and physical exertion. For stress, there was a significant main effect of environment, with higher ratings in the gray compared to the green environment (*b* = 7.81, *SE* = 2.23, t(73.80) = 3.50, *p* < .001). For affective well-being, participants reported significantly lower ratings in gray compared to green (*b* = -6.28, *SE* = 2.40, *t*(77.18) = -2.61, *p* = .011). Time, order group, and their interactions with environment were non-significant across both models (all *p* > .14; see Table S2 for full results). Overall, exposure to green environments was associated with reduced subjective stress and improved affective well-being, independent of order or repeated exposures, while perceived physical exertion was not significantly different across conditions.

**Figure 3.**
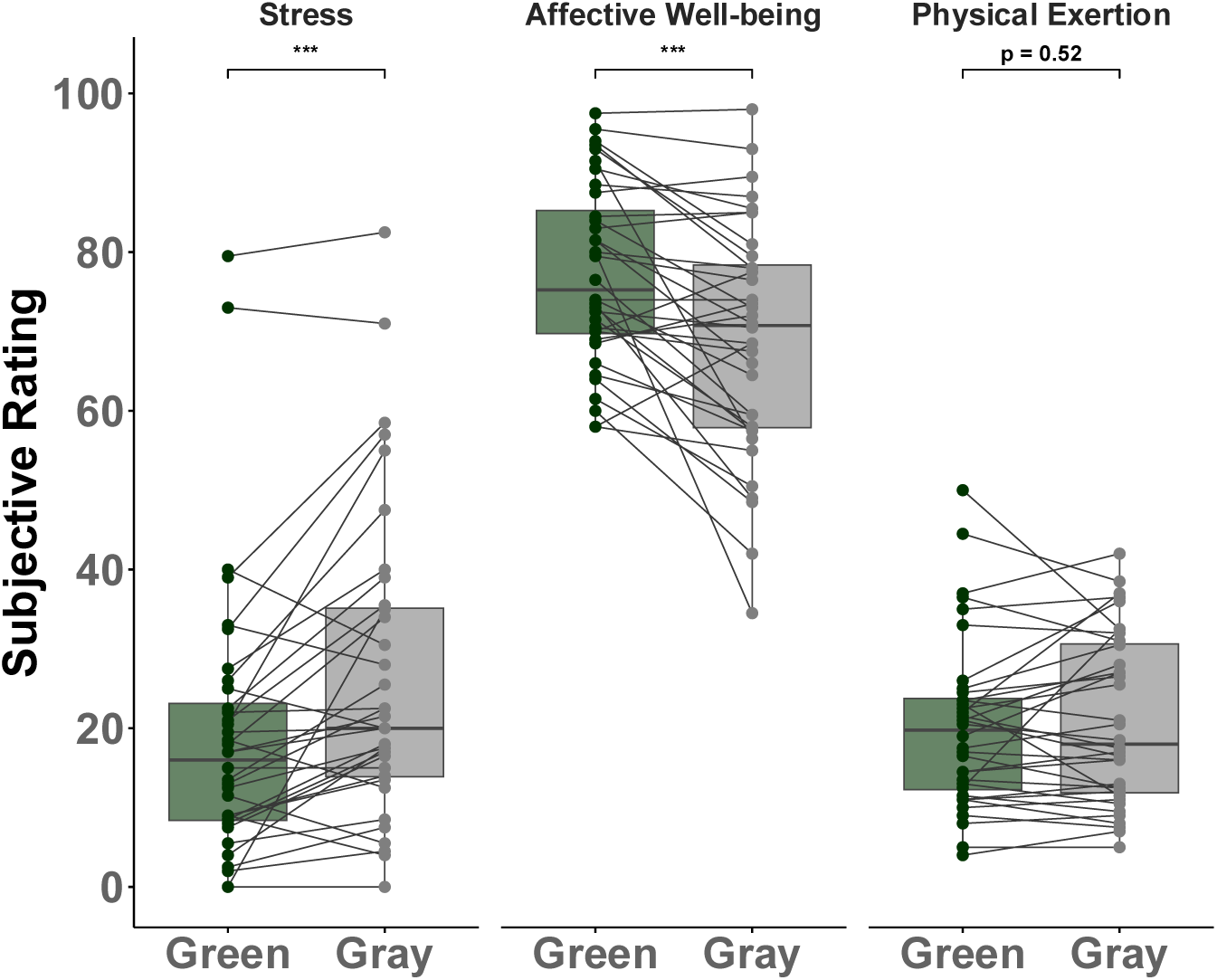
Paired comparisons of subjective ratings between green and gray urban environments for three EMA outcomes: stress, affective well-being, and physical exertion (all rated 0–100). Individual subject trajectories are connected by lines; boxes represent the interquartile range with the median. One missing gray (*n*=1) EMA data. Significance brackets reflect paired t-tests (* *p* < .05, ** *p* < .01, *** *p* < .001). *N*= 36.

Pearson correlations amongst the different EMA ratings differed numerically across environments. In the green environment, only affective well-being and physical exertion were significantly correlated (*r* = -.37, *p* < .05), whereas stress showed no significant associations with the other items. In the gray environment, stress was significantly associated with both affective well-being (*r* = -.33, *p* < .05) and physical exertion (r = .35, *p* < .05), while the correlation between affective well-being and exertion did not reach significance (see Fig. S2).

### Physiology

Visual inspection of route-based physiological trajectories revealed a pronounced increase in HR in one green segment, attributable to a notable incline slope confirmed by on-site route verification. This segment was excluded from subsequent analyses on the basis of a significant HR elevation relative to the unaffected green segment (Supplementary Fig. S3). Additionally, preregistered LMMs and correlation analyses examining the relationship between physiological indices and subjective ratings (see Table S3; Fig. S2), as well as an assessment outside temperature (see Table S4), are reported in the *supplementary information*.

#### Segment-based analyses

Paired-samples t-tests with Bonferroni correction across five physiological variables (α = .01) revealed no significant differences between green and gray environments in the full analysis (all *p* > .05, Fig. 4a). When the green segment confounded by the slope was excluded (see Fig. 4b), mean HR was significantly lower during green (*M* = 97.61, *SD* = 11.19) compared to gray space exposure (*M* = 99.69, *SD* = 11.25; *t*(35) = −4.31, *p* < .001, *d* = −0.72). Heart rate variability showed a trend-level effect (*p* = 0.054).

**Figure 4.**
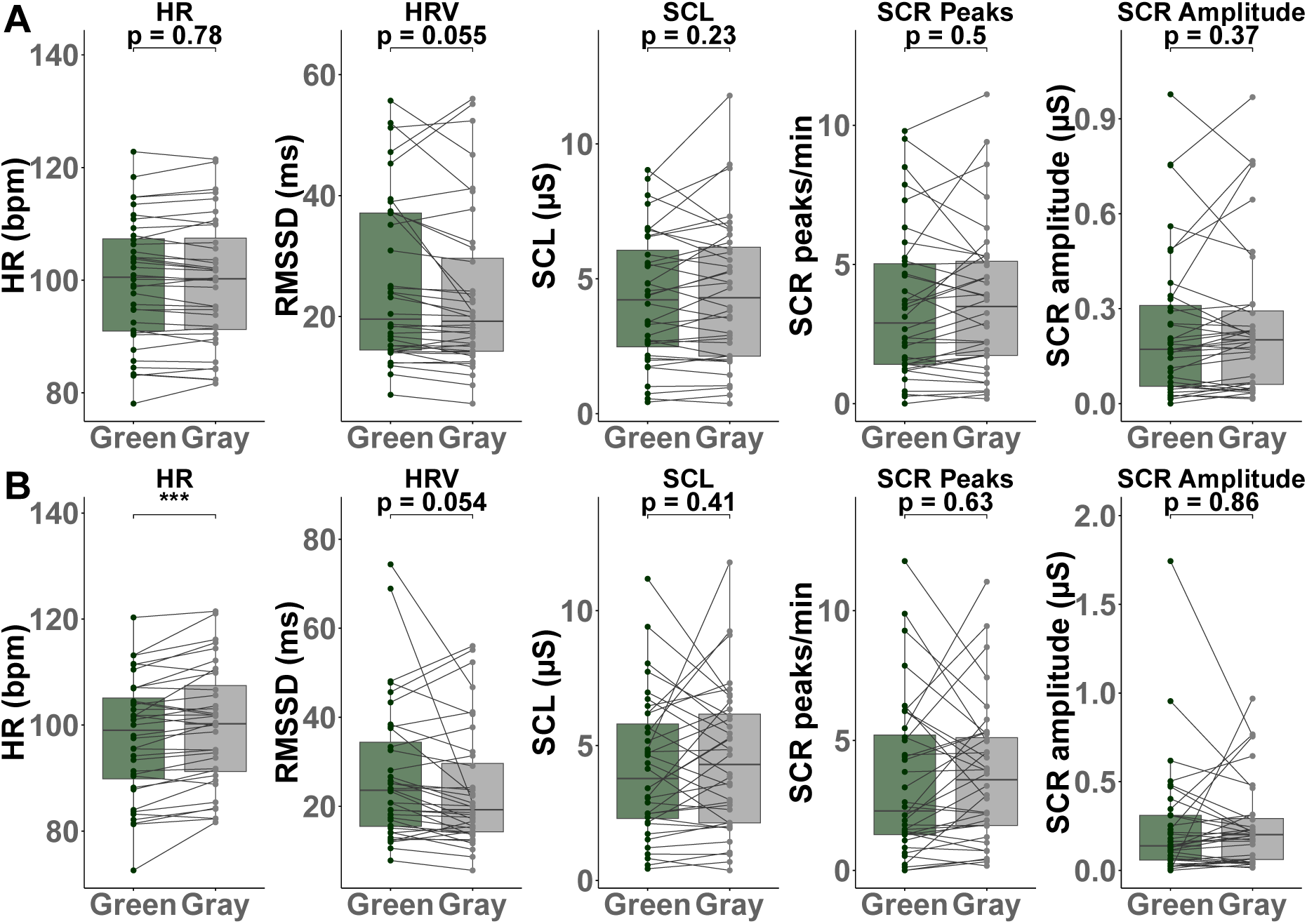
Paired physiological responses across green and gray urban environments for five measures: heart rate (HR, bpm), heart rate variability (HRV, RMSSD in ms), skin conductance level (SCL, µS), skin conductance response peak rate (SCR, peaks/min), and SCR amplitude (µS). Panel A shows the full analysis including all green segments; panel B shows the slope-corrected analysis with the confounded green segment excluded. Individual subject trajectories are connected by lines; boxes represent the interquartile range with the median; significance brackets indicate Bonferroni-corrected paired t-test results (* *p* < .05, ** *p* < .01, *** *p* < .001). *N* = 36.

We next ran a PCA analysis to take the covariance across physiological measures into account (Fig. 5). In the full analysis, PC1 explained 46.4% of variance and was characterized by high negative loadings on EDA variables, reflecting (reduced) electrodermal arousal. PC2 explained 26.9% of variance and was dominated by cardiac variables, with a strong negative loading on HR and a positive loading on heart rate variability, reflecting cardiac recovery. Together, PC1 and PC2 accounted for 73.3% of total variance (see Fig. 5a). In the slope-excluding analysis, the factor structure was largely preserved (Fig. 5b). PC1 again reflected electrodermal arousal (49.9% variance) and PC2 again reflected cardiac arousal (23.4% variance). Together these two components accounted for 73.3% of variance, consistent with the full analysis, suggesting that the slope exclusion did not substantially alter the underlying correlation structure (see Fig. 5b).

**Figure 5.**
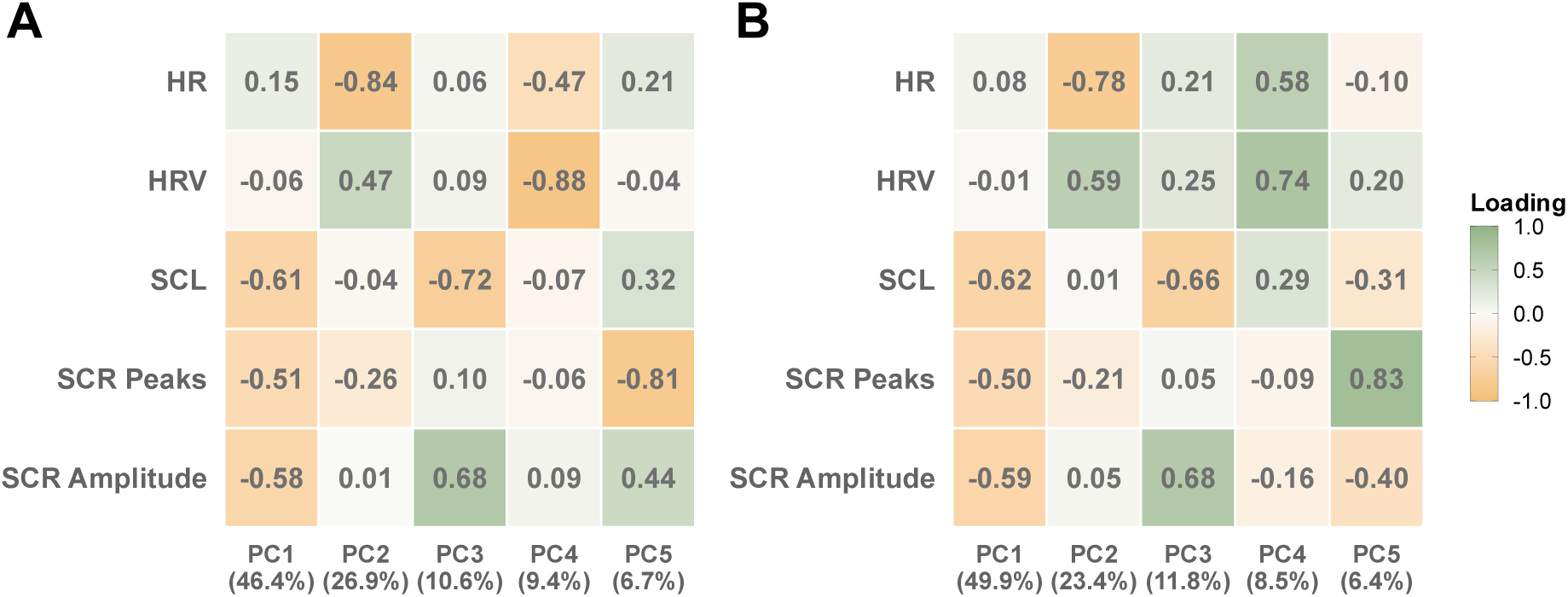
Heatmap of PCA factor loadings and explained variance for the segment-based physiological analysis, separately for the full analysis (panel A) and the analysis with the confounded green segment excluded (panel B). Rows represent five physiological variables: heart rate (HR, bpm), heart rate variability (HRV; RMSSD, ms), skin conductance level (SCL, µS), SCR peaks per minute, and SCR amplitude (µS). Columns represent principal components (PC1–PC5) with explained variance in parentheses. PCA was conducted on within-subject *z*-scored physiological variables. *N* = 36.

Paired t-tests on aggregated PC scores revealed no significant differences between green and gray environments in the full analysis (all *p* > .07, see Fig. 6a). In the slope-excluding analysis, the electrodermal component (PC1) did not differ significantly between environments (*t*(35) = 0.76, *p* = .452, *d* = 0.13). However, the cardiac component (PC2) was significantly increased during green (*M* = 0.39, *SD* =.81) compared to gray exposure (*M* = −0.19, *SD* = .41, *t*(35) = 3.41, *p* = .002, *d* = 0.57), reflecting reduced HR and increased heart rate variability during green vs. gray space exposure. All remaining components were non-significant (all *p* > .57).

**Figure 6.**
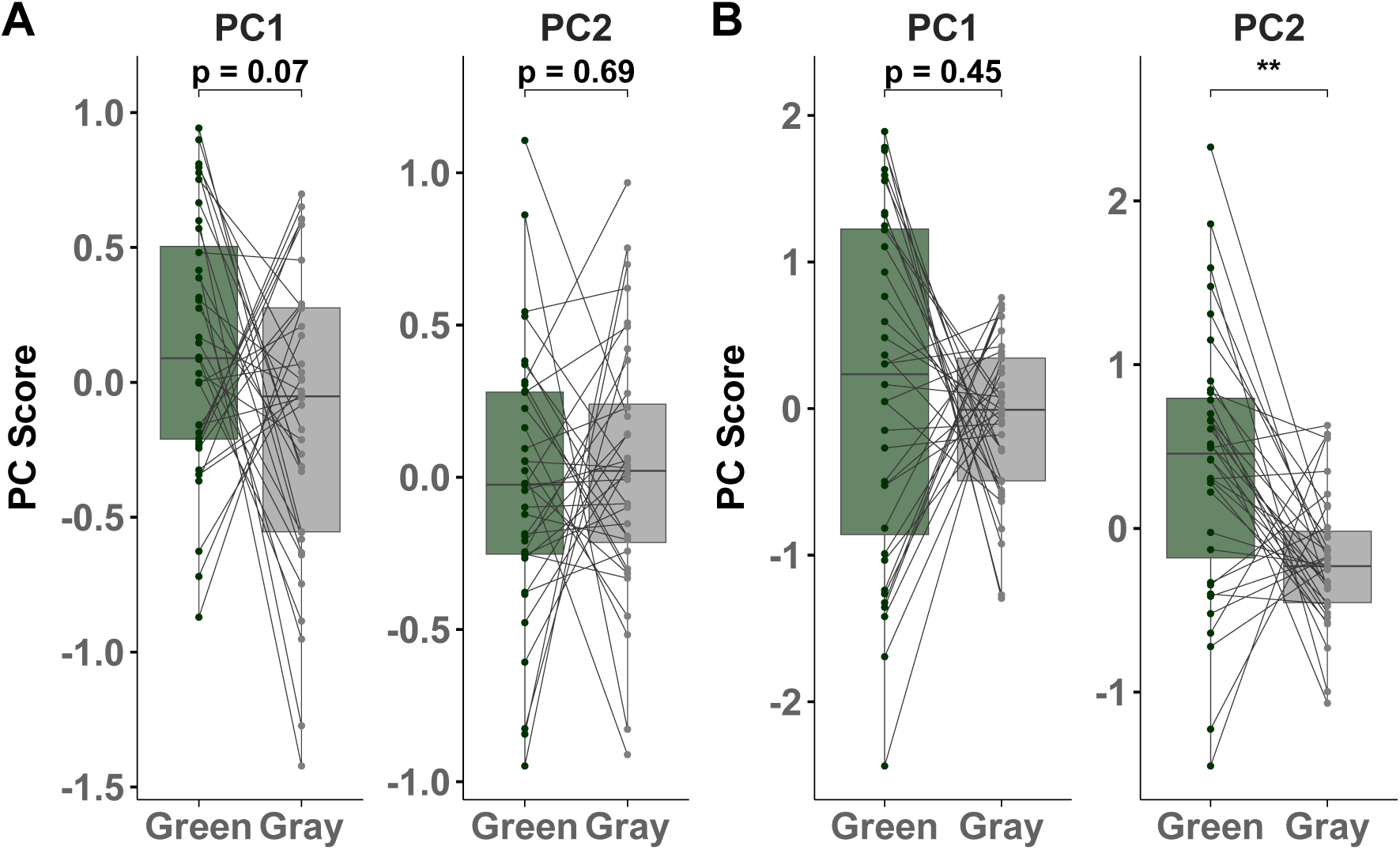
PCA scores comparing green and gray urban environments across two principal components (PC1–PC2). Panel A shows the full analysis including all green segments; panel B shows the slope-corrected analysis with the confounded green segment excluded. Single-subject trajectories are connected by lines; boxes represent the interquartile range with the median. PC scores are derived from within-subject z-scored physiological variables: heart rate (HR, bpm), heart rate variability (HRV, RMSSD in ms), skin conductance level (SCL, µS), skin conductance response peak rate (SCR, peaks/min), and SCR amplitude (µS). Significance brackets reflect paired t-tests (* *p* < .05, ** *p* < .01). *N* = 36.

#### Transition-based analysis

PCA results for the transition-based analysis were highly similar to those of the segment-based analysis, and are summarized in *supplementary information* (see Fig. S4). The bin-wise PC score trajectories depicting the temporal dynamics of autonomic responding across the nine-minute transition window are shown in Figure 7, separately for each transition direction and for both the full and slope-excluded analyses. In the full analysis, PC score trajectories showed considerable variability around the moment of transition across both directions. A notable downward shift in PC2 was observed at Bin 0 for both transition directions, indicating a transient increase in cardiac arousal at the moment of environmental change regardless of direction.

**Figure 7.**
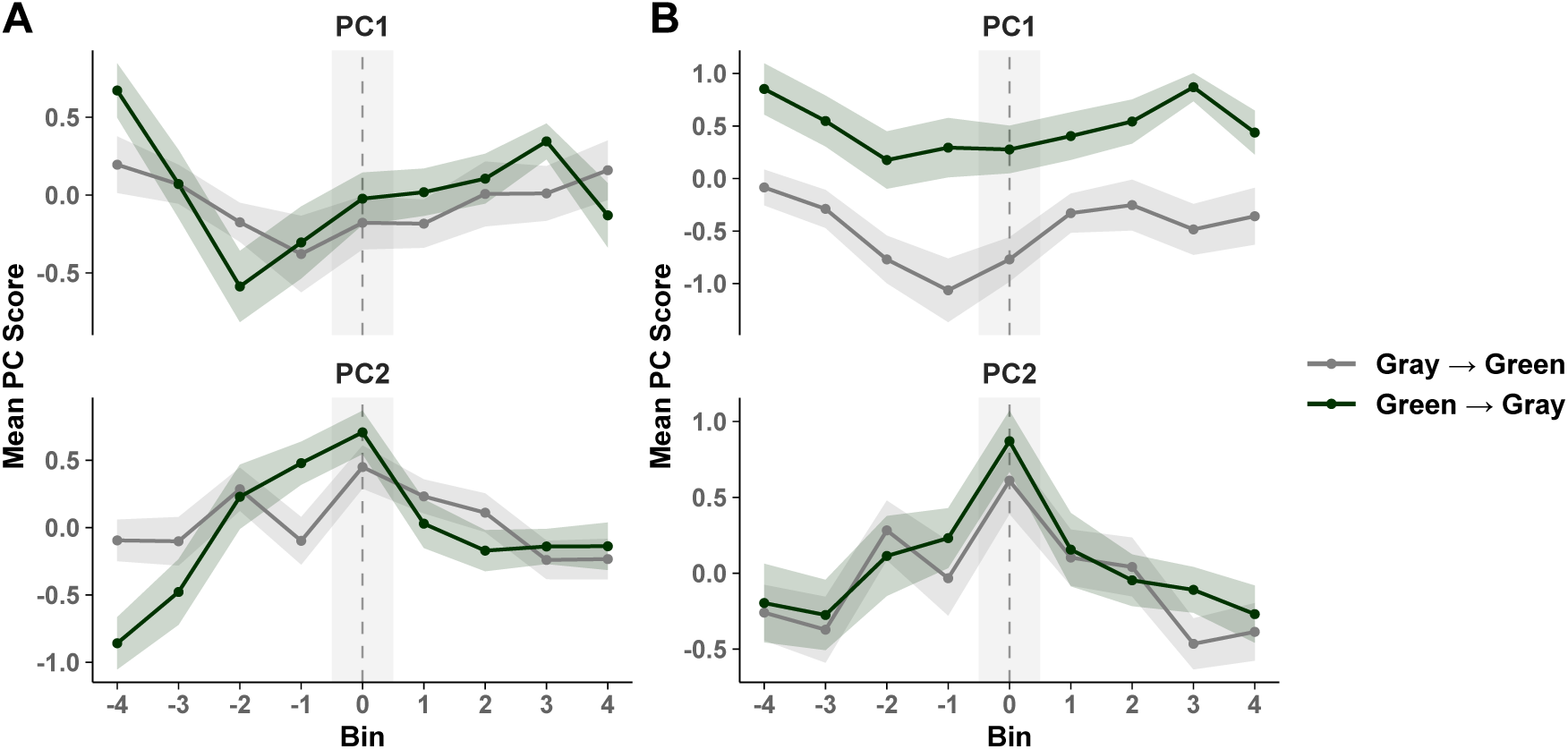
PCA-derived PC score time-courses across nine 60-second bins relative to the moment of environmental transition (0 = transition) for green-to-gray (dark green) and gray-to-green (gray) transitions. Panel A depicts all transitions included; panel B shows slope-corrected transitions excluded. Negative bins indicate the pre-transition period; positive bins indicate the post-transition period. Shaded ribbons indicate ± 1 standard error of the mean; the gray shaded band marks the transition bin.

The slope-excluded analysis revealed more interpretable and directionally distinct patterns (see Fig. 7b). For gray-to-green transitions, PC1 scores (representing electrodermal arousal, where lower values indicate higher activation) were most negative in the pre-transition period, reflecting peak electrodermal arousal during the gray environment. Following entry into the green segment, these scores showed a gradual positive shift, reflecting a reduction in electrodermal arousal. For green-to-gray transitions, PC1 scores were positive prior to the transition, reflecting reduced electrodermal arousal during the green segment. PC1 scores remained elevated even after re-entry into the gray environment, indicating a possible carry-over of parasympathetic dominance from the green into the subsequent gray segment. In PC2 (representing cardiac activity, where positive values indicate higher vagal activity), both transition directions showed a prominent transient increase at bin 0. This reflects a brief decrease in HR and a corresponding increase in heart rate variability at the exact moment of environmental crossing.

## Discussion

The present study examined the impact of urban green versus gray space exposure on subjective stress levels and physiological arousal during naturalistic urban mobility. Participants reported lower subjective stress and higher affective well-being in green compared to gray environments, with moderate to large effect sizes, while cardiac measures indicated reduced HR and elevated heart rate variability consistent with a shift toward parasympathetic dominance during green space exposure. Transition-based analyses further indicate arousal carry-over effects persisting beyond immediate environmental boundaries, suggesting that even brief green space encounters during everyday urban mobility are sufficient to elicit stress reduction at both psychological and autonomic levels.

Subjective stress and affective well-being differed significantly between green and gray segments, with effect sizes (*d* = 0.66 and *d* = 0.71) comparable to or exceeding those reported in laboratory and virtual reality studies (62,63), likely reflecting the richer multisensory character of naturalistic exposure (33,39). The absence of differences in perceived physical exertion confirms that these effects reflect genuine environmental influences rather than differential effort. Critically, effects were robust across repeated exposures, with neither order nor time effects observed, consistent with *Stress Recovery Theory’s* proposal that nature-elicited recovery is rapidly driven by environmental affordances (24), and with *Attention Restoration Theory’s* prediction that even brief encounters with natural settings replenish depleted cognitive resources (25). Prior evidence has linked longer or cumulative nature exposure to improved well-being (35,64); the present findings suggest that comparable restorative processes operate during much shorter exposures embedded in everyday urban mobility, indicating that the benefits of green space may extend across a broader temporal spectrum than previously assumed.

At the physiological level, initial segment-based analyses did not yield significant electrodermal or cardiac differences, but correcting for the slope-affected segment revealed significantly lower HR in green compared to gray environments. A PCA confirmed this pattern, identifying a cardiac component (PC2) loading negatively on HR and positively on heart rate variability, whose scores were significantly reduced during green space exposure, indicating lower physiological arousal and replicating laboratory and field findings (24,34,37,40), under naturalistic conditions with repeated environmental transitions. The absence of significant segment-level electrodermal differences likely reflects the distinct functional properties of EDA: tonic SCL indexes reflect slow sympathetic buildup requiring sustained or intense exposure to differentiate conditions, while phasic SCR peaks respond to discrete stimuli rather than sustained environmental states (10,11,65). Attenuated tonic reactivity among urban residents habituated to these environments (10,66), compounded by cold autumn and winter temperatures dampening sweat gland activity (10) may have further limited segment-level electrodermal differentiation.

Transition-based analyses centered on GPS-verified environmental boundaries revealed dynamics absent from segment-level aggregates, underscoring the sensitivity of ambulatory assessments to effort confounds and the importance of careful data quality control. The electrodermal component (PC1) showed a directional asymmetry across transition types. During gray-to-green transitions, electrodermal arousal was markedly elevated in the pre-transition gray period and declined progressively following green space entry, while during green-to-gray transitions, suppressed arousal persisted well into the subsequent gray segment. This autonomic carry-over effect extends previous evidence of post-exposure subjective well-being benefits (42) to the physiological level and aligns with *Stress Recovery Theory’s* prediction of a sustained recovery trajectory following nature exposure (24), suggesting that brief green space encounters may partially buffer against the arousal-inducing properties of subsequent gray environments. The cardiac component (PC2) exhibited a consistent pattern across both transition directions, with a brief non-directional peak at the transition point likely reflecting an orienting response to environmental novelty or anticipatory effects. The dissociation between this transient spike and the sustained directional environmental effect suggests that the cardiac component may capture two distinct processes, highlighting the importance of examining multiple temporal dynamics when interpreting autonomic responses in naturalistic settings.

Green and gray segments could not be perfectly matched due to geographic constraints, with gray segments approximately twice as long in duration and distance (*M*_gray_ = 16.18 min, 1064m; *M*_green_ = 9.22 min, 652m). That restorative effects emerged despite participants spending less time and walking faster through green than gray environments suggests that environmental quality rather than exposure duration drives recovery (24,67), though future studies should aim for closer matching. Effects were further observed across a wide range of seasonal and meteorological conditions, including temperatures from below freezing to 20°C and varying vegetation cover, positioning urban green infrastructure as a resilient stress-regulatory resource even outside optimal conditions, consistent with large-scale evidence supporting nature-based interventions across diverse contexts (22,24). Whether seasonal factors systematically moderate physiological responses warrants further investigation.

Subjective and physiological measures were not significantly correlated within either environment, and physiological indices did not predict momentary subjective stress, consistent with theoretical accounts of partial independence between psychological and autonomic stress systems (24) and with evidence that self-reported and physiological measures may diverge in naturalistic settings (68). This may also reflect temporal mismatch between discrete EMA assessments and continuously recorded signals. Importantly, this dissociation did not prevent green environments from producing consistent effects across both levels, suggesting that green space exposure may affect multiple response systems via partially independent pathways, with cognitive and attentional mechanisms operating rapidly as proposed by *Attention Restoration Theory* (25), and autonomic recovery unfolding more gradually through the stress-regulatory processes described in *Stress Recovery Theory* (24). These findings underscore the importance of jointly examining subjective and physiological measures alongside their distinct temporal profiles (69–71).

### Limitations

Several limitations should be considered when interpreting the findings. The sample consisted of healthy young students, and whether effects extend to other demographic groups (72) or individuals with stress-related mental health conditions remains to be investigated. All participants navigated the same fixed route within a single city, and future studies employing multiple routes or diverse green space types could reveal specific contributing environmental factors and enhance generalizability. Physiological assessment covered cardiac and electrodermal indices but not HPA measures such as salivary cortisol, where dysregulation is implicated in mental disorders and health outcomes (73,74). Future studies should adapt designs to capture neuroendocrine responses alongside autonomic indices. Geographic constraints precluded perfect matching of green and gray segment lengths and introduced a topographic confound in one segment, both of which future studies should address through closer environmental matching and improved route consistency. Finally, data collection was restricted to autumn and winter in a temperate climate, and systematic investigation across seasons and varying weather conditions would clarify the extent to which the present findings generalize across the year.

## Conclusion

In conclusion, the present study provides multi-level evidence that brief and repeated contact with urban green space supports stress recovery during urban mobility in a setting with high ecological validity. Green environments were consistently associated with reduced subjective stress, higher affective well-being, and reduced cardiac autonomic arousal across repeated environmental transitions, with physiological effects persisting beyond the immediate exposure window. Together, these findings underscore the public health potential of urban green infrastructure as a simple and scalable strategy for stress regulation and the prevention of stress-related mental health conditions in urban populations.

## Author Contributions

Conceptualization: J.P & D.K

Data curation: D.K

Formal analysis: D.K

Funding acquisition: J.P

Investigation: Q.B, M.M, & D.K

Methodology: J.P & D.K

Project administration: J.P & D.K

Resources: J.P

Software: RStudio 2024 & Python

Supervision: J.P

Validation: J.P & D.K

Visualization: D.K

Writing – original draft: D.K

Writing – review & editing: J.K & J.P

## Acknowledgements

This work was funded by the German Research Foundation (DFG, grant 502778657 to J.P.). J.K. and J.P. further acknowledge financial support from the Mapping Autonomic Neural Interaction and Control (MANIAC) Emerging Group by the University of Cologne Excellent Research Support Program via the German Research Foundation (DFG).

## Supplementary Information

### Supplementary Methods

#### Measures

##### Stress Overload Scale

The Stress Overload Scale short version (75) assesses stress overload across two subscales. Participants reported moderate levels of stress overload for the event load subscale (*M(SD) =* 16.06 (5.04); range = 6–23) and the personal vulnerability subscale (*M(SD)* = 11.33 (3.97); range = 5–22). The total score had a mean of *M(SD) =* 27.39 (7.81); range = 13–45). Internal consistency was high for the total scale (α = .85) and the event load subscale (α = .88), and acceptable for the personal vulnerability subscale (α = .72).

##### Perceived Stress Scale

The Perceived Stress Scale short version (PSS-10) (76) measures perceived stress across two subscales. The perceived helplessness subscale (*M(SD)* = 17.11 (3.41); range = 10–25), and the perceived self-efficacy subscale *(M(SD)* =10.56 (2.60); range = 6–16). The total PSS-10 score had a mean of *M(SD)* = 30.56 (3.09) (range = 23–36). Cronbach’s alpha indicated acceptable internal consistency for the total scale (α = .77), with good reliability for the perceived helplessness (α = .69) and perceived self-efficacy (α = .67) subscales.

##### Scale of Positive and Negative Experience

The Scale of Positive and Negative Experience (SPANE)(77) assesses affective experience across two subscales. Positive experiences (SPANE-P) were rated *M(SD)* = 21.92 (3.61; range = 10–29) and negative experiences (SPANE-N) M(SD) = 15.44 (3.48; range = 8–24). The balanced affect score (SPANE-B) was M(SD) = 6.47 (5.82; range = −14 to 16). Internal consistency was good to high across subscales (SPANE-P: α = .86; SPANE-N: α = .68; SPANE-B: α = .80).

##### State-Trait Anxiety-Inventory

The State-Trait Anxiety Inventory short version (STAI) (78) assesses both dispositional and situational anxiety. Trait anxiety was *M(SD)* = 39 (9.23; range = 22–63) and state anxiety *M(SD)* = 33.94 (11.06; range = 13–61). Internal consistency was acceptable to good across subscales (Trait: α = .70; State: α = .84).

##### Big Five Inventory

Big Five Inventory 10-Item Version (BFI-10) (79) measures the five major personality dimensions. Scores were highest for Openness *M(SD)* = 3.78 (0.97; range = 1.5–5.0) and Conscientiousness *M(SD)* = 3.62 (0.83; range = 2.0–5.0), followed by Extraversion *M(SD)* = 3.49 (0.91; range = 1.5–5.0), Agreeableness *M(SD)* = 3.39 (0.98; range = 1.5–5.0), and Neuroticism *M(SD)* = 3.19 (0.96; range = 1.0–4.5). Internal consistency ranged from acceptable to moderate across dimensions (Extraversion: α = .78; Agreeableness: α = .75; Openness: α = .65; Neuroticism: α = .62), with lower consistency for Conscientiousness (α = .47), which is expected given the two-item structure of each dimension.

##### The Satisfaction with Life Scale

The Satisfaction with Life Scale (SWLS) (80) measures cognitive life satisfaction. Total scores were *M(SD)* = 18.92 (3.17; range = 14–25), corresponding to a mean item score of *M(SD)* = 3.78(0.63; range = 2.8–5.0), indicating moderate life satisfaction. Internal consistency was acceptable (α = .69), consistent with the brevity of the five-item scale.

##### Brief Scale Sensation Seeking

The brief Sensation Seeking Scale (BSSS-4) (81) assesses sensation seeking with four items. Total scores were *M(SD)* = 11.17 (3.68; range = 5–19), indicating moderate sensation seeking. Internal consistency was good (α = .78), consistent with previous reports for this scale.

##### Connectedness to Nature

The Connectedness to Nature Scale (CNS) (82) measures the degree to which individuals feel connected to the natural world. Total scores were *M(SD)* = 42.14 (8.16; range = 21–60), indicating moderate to relatively high nature connectedness. Internal consistency was good (α = .80), supporting the reliability of the translated version used in the present study.

## Supplementary Tables

**Table S1.** Descriptive Statistics for Ecological Momentary Assessment Ratings by Environment Segment. Means (*M*), standard deviations (*SD*), and observed ranges are reported for momentary stress, affective well-being, and physical exertion, each assessed on a 0–100 visual analogue scale. The number of ecological momentary assessments (EMA) prompts per segment reflects the study protocol: single assessments were administered at the baseline and post-exposure segments, and two assessments were administered within each green and gray exposure segment. Minor deviations from the expected prompt count (*n* = 36 per segment) reflect isolated missing responses in the baseline (*n* = 2), gray (*n* = 1), and post-exposure (*n* = 1) segments. Green and gray values represent aggregated means across the two repeated exposures within each environmental condition. *N* = 36.

| Environment Segment | EMA prompts | Stress |  | Affective well-being |  | Physical Exertion |  |
| --- | --- | --- | --- | --- | --- | --- | --- |
|  |  | *M* (*SD*) | Range | *M* (*SD*) | Range | *M* (*SD*) | Range |
| Baseline | 34 | 24.36 (24.02) | 0-91 | 72.39 (14.19) | 38-100 | 10.39 (7.47) | 0-31 |
| Gray | 71 | 26.03 (19.76) | 0-82.5 | 69.18 (14.73) | 45.5-98 | 20.96 (10.68) | 5-42 |
| Green | 72 | 18.94 (17.64) | 0-79.5 | 77.24 (11.33) | 58-97.5 | 20.24 (10.66) | 4-50 |
| Post | 35 | 21.33 (19.03) | 0-79 | 77.33 (14.81) | 49-100 | 22.27 (15.28) | 4-60 |

**Table S2.** Linear Mixed-Effects Models Predicting Subjective Stress, Affective Well-being, and Perceived Exertion from Green Space Exposure. Subjective stress, affective well-being, and perceived exertion assessed via EMA served as outcome variables. Environment was dummy-coded (Gray = 0, Green = 1), such that positive *B* values indicate higher scores in green vs. gray environments. Measurement number (1 = first prompt, 2 = second prompt within each environmental segment) was included to test for within-segment changes over time. Group order reflects the counterbalanced assignment to green-first or gray-first walking sequences. Values represent unstandardized regression coefficients (*B*). All models included a random intercept for participant to account for the nested structure of observations within participants. *N* = 36. \**p* < .05.

| Variable | Predictor | *B* | SE | df | *t* | *p* |
| --- | --- | --- | --- | --- | --- | --- |
| Stress | Intercept | 18.49 | 4.31 | 37.67 | 4.29 | < .001 |
|  | Environment (Gray vs Green) | 7.81 | 2.23 | 73.80 | 3.50 | **.001** |
|  | Measurement | -1.28 | 1.89 | 70.00 | -0.68 | .502 |
|  | Group Order | 2.19 | 5.93 | 34.00 | 0.37 | .715 |
|  | Environment × Measurement | -1.44 | 2.68 | 70.00 | -0.54 | .591 |
| Affect | Intercept | 76.34 | 2.89 | 45.28 | 26.43 | < .001 |
|  | Environment (Gray vs Green) | -6.28 | 2.40 | 77.18 | -2.61 | **.011** |
|  | Measurement | 3.08 | 2.11 | 70.00 | 1.47 | .148 |
|  | Group Order | -1.28 | 3.78 | 34.00 | -0.34 | .736 |
|  | Environment × Measurement | -3.56 | 2.98 | 70.00 | -1.19 | .236 |
| Exertion | Intercept | 19.50 | 2.64 | 44.78 | 7.40 | < .001 |
|  | Environment (Gray vs Green) | -0.72 | 1.75 | 104.95 | -0.41 | .564 |
|  | Measurement | 3.14 | 1.75 | 104.99 | 1.79 | .076 |
|  | Group Order | -1.66 | 3.42 | 34.00 | -0.49 | .630 |
|  | Environment × Measurement | 2.89 | 2.48 | 104.99 | 1.17 | .246 |

**Table S3.** Linear Mixed-Effects Models Predicting Subjective Stress from Green Space Exposure and Physiological Indices. Values represent unstandardized regression coefficients (*B*). Subjective stress assessed via EMA served as the outcome variable. Environment was dummy-coded (Gray = 0, Green = 1), such that positive *B* values indicate higher stress in green vs. gray environments. Each physiological index was entered separately as a predictor alongside environment: heart rate (HR, bpm), heart rate variability (HRV; RMSSD, ms), skin conductance level (SCL, µS), skin conductance response (SCR) peaks per minute, and SCR amplitude (µS). All physiological indices reflect within-subject standardized *z*-scores. All models included a random intercept for participant to account for the nested structure of observations within participants. *N* = 36. \**p* < .05.

| Variable | Predictor | *B* | *SE* | df | *t* | *p* |
| --- | --- | --- | --- | --- | --- | --- |
| Stress | Intercept | 26.00 | 03.04 | 40.80 | 8.55 | < .001 |
|  | Environment (Gray vs Green) | -7.07 | 1.51 | 108.00 | -4.69 | < .001 |
|  | HR | -0.06 | 0.87 | 108.00 | -0.07 | .941 |
|  | Intercept | 26.10 | 03.04 | 40.80 | 8.57 | < .001 |
|  | Environment (Gray vs Green) | -7.22 | 1.51 | 108.00 | -4.78 | < .001 |
|  | HRV | 0.50 | 0.87 | 108.00 | 0.57 | .567 |
|  | Intercept | 26.20 | 03.04 | 40.70 | 8.61 | < .001 |
|  | Environment (Gray vs Green) | -7.43 | 1.49 | 108.00 | -4.99 | < .001 |
|  | SCL | -1.48 | 0.86 | 108.00 | -1.73 | .087 |
|  | Intercept | 26.10 | 03.04 | 40.50 | 8.60 | < .001 |
|  | Environment (Gray vs Green) | -7.30 | 1.47 | 108.00 | -4.98 | < .001 |
|  | SCR Peaks | -1.85 | 0.85 | 108.00 | -2.19 | **.031\*** |
|  | Intercept | 26.10 | 03.04 | 40.70 | 8.59 | < .001 |
|  | Environment (Gray vs Green) | -7.28 | 1.50 | 108.00 | -4.86 | < .001 |
|  | SCR Amplitude | -0.89 | 0.87 | 108.00 | -1.03 | .307 |

**Table S4.** Correlations between Temperature, Ecological Momentary Assessment, and Physiological Indices. Outdoor temperature was recorded at the time of each measurement. Physiological indices included heart rate (HR, bpm), heart rate variability (HRV; RMSSD, ms), skin conductance level (SCL, µS), skin conductance response (SCR) peaks per minute, and SCR amplitude (µS). Values represent Pearson’s *r*. All physiological variables were standardized within subjects prior to analysis. Significant correlations after Bonferroni correction are marked with an asterisk (\**p* < .05.). *N* = 36.

| Variable | 1 | 2 | 3 | 4 | 5 | 6 | 7 | 8 | 9 |
| --- | --- | --- | --- | --- | --- | --- | --- | --- | --- |
| 1. Temperature | - |  |  |  |  |  |  |  |  |
| 2. Stress | -.11 | - |  |  |  |  |  |  |  |
| 3. Affective Well-being | .08 | -.04 | - |  |  |  |  |  |  |
| 4. Physical Exertion | -.12 | .28 | **-.37\*** | - |  |  |  |  |  |
| 5. HR | -.12 | -.10 | .09 | -.04 | - |  |  |  |  |
| 6. HRV | .08 | -.07 | .07 | .00 | **-.53\*** | - |  |  |  |
| 7. SCL | -.14 | .07 | -.06 | .19 | .05 | .03 | - |  |  |
| 8. SCR Peaks | -.29 | .18 | -.02 | .10 | .32 | -.14 | **.52\*** | - |  |
| 9. SCR Amplitude | -.26 | .09 | -.04 | -.05 | -.09 | -.13 | **.43\*** | **.51\*** | - |

## Supplementary Figures

**Figure S1.**
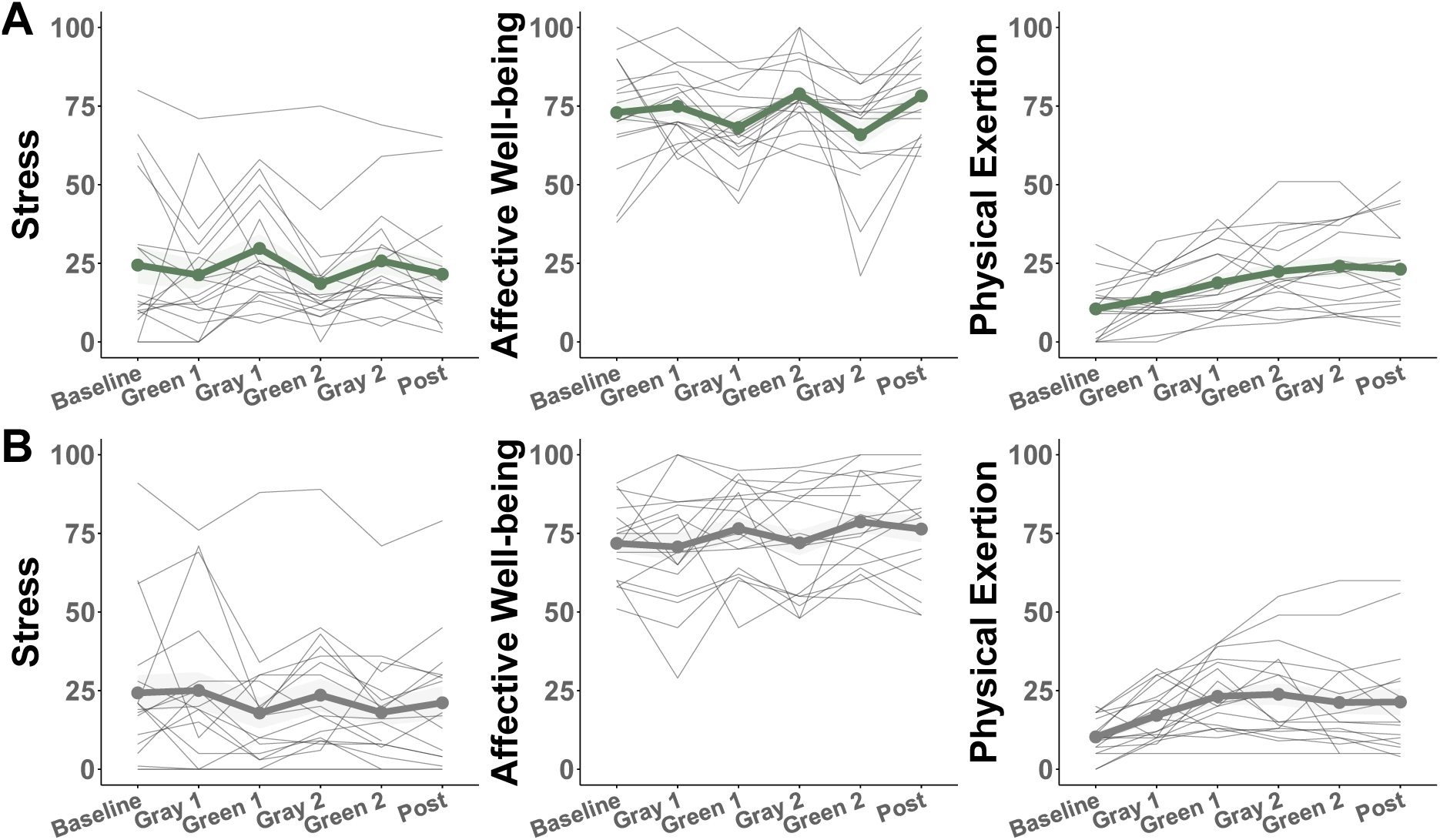
Ecological Momentary Assessment Trajectories across Segments. Ecological momentary assessment trajectories across six measurement points (Baseline, Green 1/Gray 1, Green 2/Gray 2, Post) for three subjective outcomes: stress, affective well-being, and physical exertion (all rated 0–100). Panel A depicts green-first participants (*n* = 18); Panel B depicts gray-first participants (*n* = 18). Thin gray lines represent individual subject trajectories; the bold line represents the group mean; shaded ribbons indicate ± 1 standard error of the mean.

**Figure S2.**
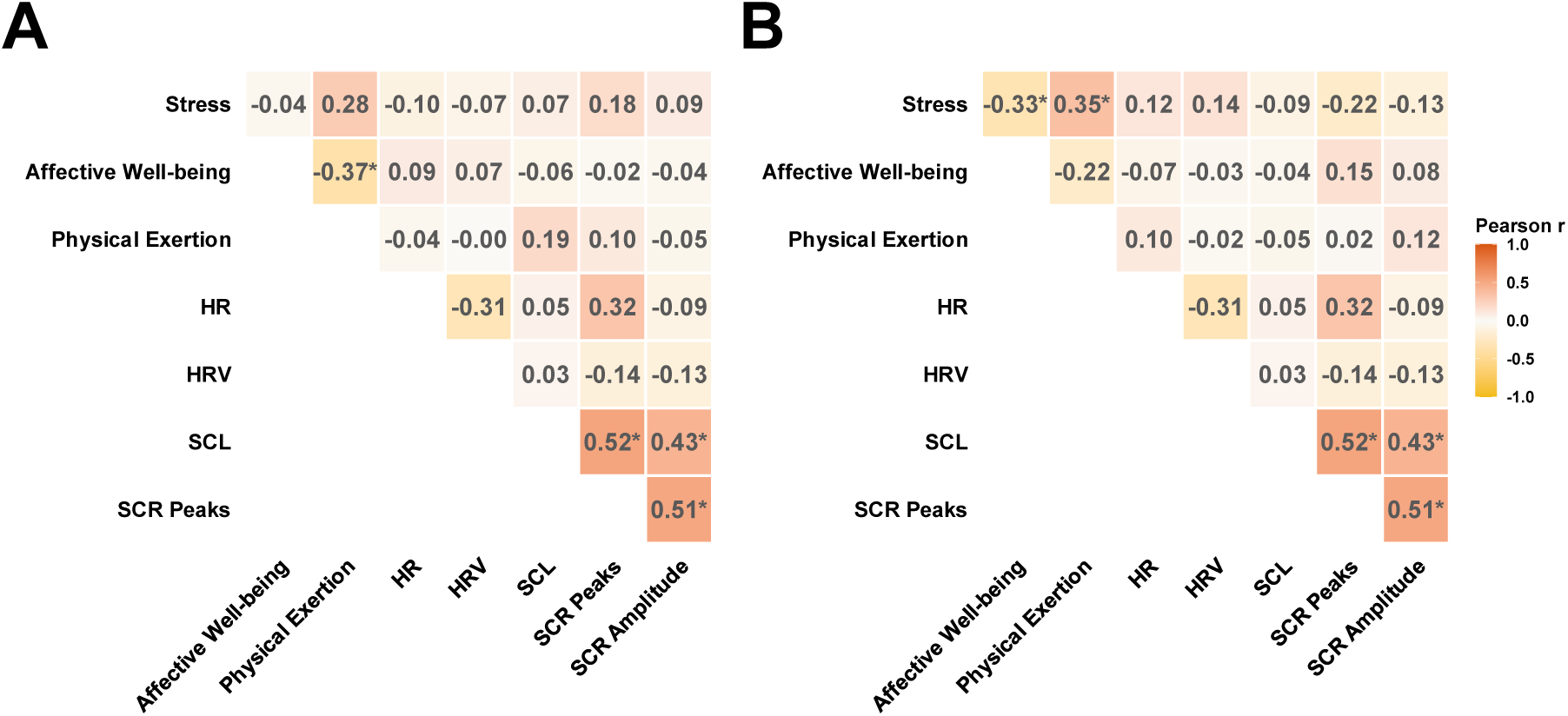
Correlation Heatmaps of Psychological and Physiological Variables across Environments. Pearson correlation coefficients (*r*) for (A) green and (B) gray environments. Variables include self-reported stress, affective well-being, and physical exertion, alongside physiological measures: heart rate (HR, bpm), heart rate variability (HRV; RMSSD, ms), skin conductance level (SCL, µS), SCR peaks per minute, and SCR amplitude (µS). Only the upper triangle of the correlation matrix is shown. \**p* < .05. *N* = 36.

**Figure S3.**
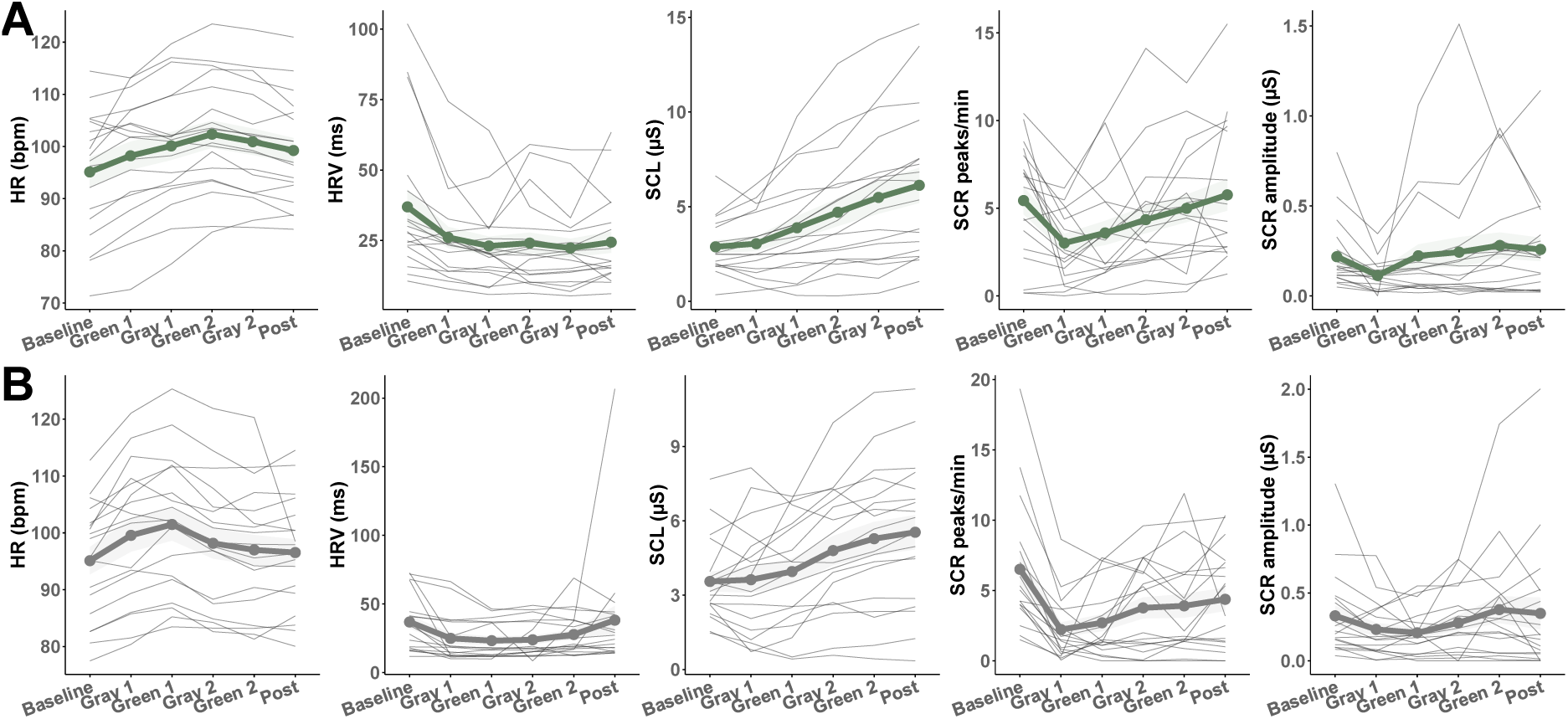
Segment-level Physiological Trajectories across Segments. Physiological trajectories across six segments (Baseline, Green 1/Gray 1, Green 2/Gray 2, Post) for five physiological measures: heart rate (HR, bpm), heart rate variability (HRV; RMSSD, ms), skin conductance level (SCL, µS), SCR peaks per minute, and SCR amplitude (µS). Panel A: green-first (n = 18); Panel B: gray-first (n = 18). Thin gray lines = individual trajectories; bold line = group mean; shaded ribbons = ±1 standard error of the mean.

**Figure S4.**
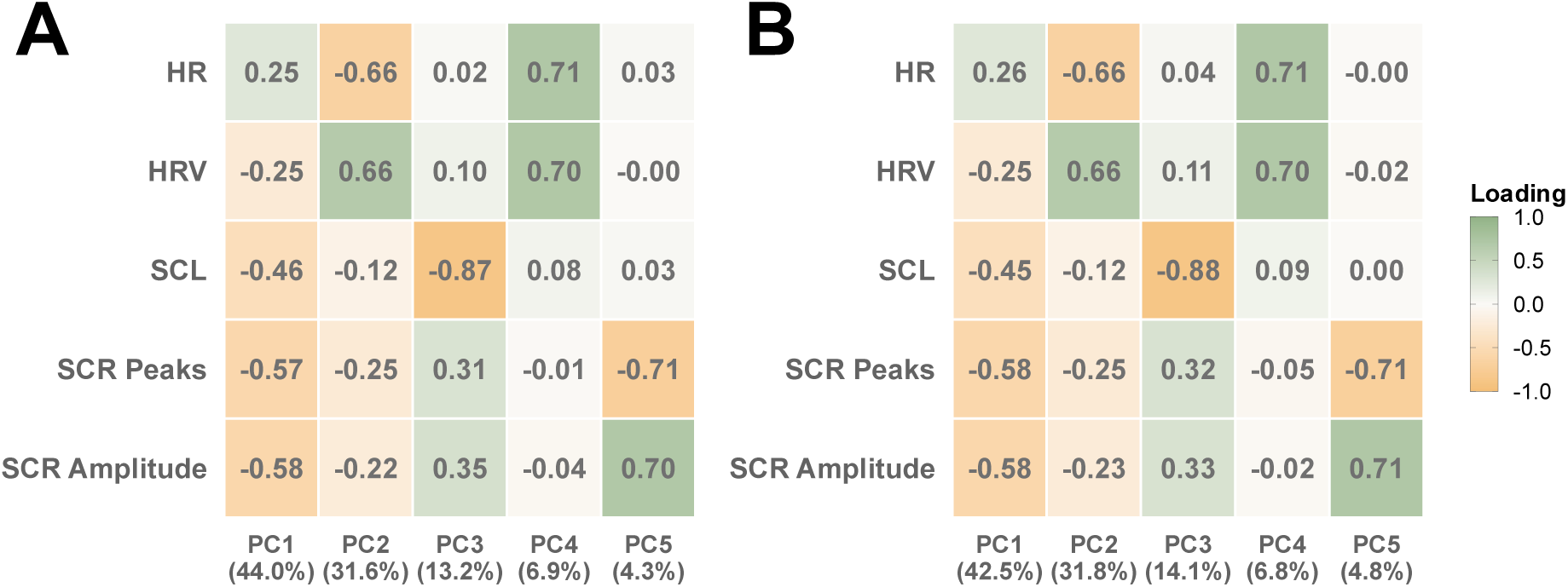
PCA Factor Loadings for Transition-based Physiological Analysis. Heatmap of PCA factor loadings and explained variance for the transition-based physiological analysis, separately for the full analysis (panel A) and the analysis with the confounded green segment excluded (panel B). Rows represent five physiological variables: heart rate (HR, bpm), heart rate variability (HRV; RMSSD, ms), skin conductance level (SCL, µS), SCR peaks per minute, and SCR amplitude (µS). Columns represent principal components (PC1–PC5) with explained variance in parentheses. PCA was conducted on within-subject *z*-scored physiological variables. *N* = 36.

## Notes

### Competing Interest Statement

The authors have declared no competing interest.

